# Non-genetic messenger RNA silencing reveals essential genes in phage-host interplay

**DOI:** 10.1101/2024.07.31.605949

**Authors:** Milan Gerovac, Svetlana Ðurica-Mitić, Leandro Buhlmann, Valentin Rech, Yan Zhu, Samuel Carien, Linda Popella, Jörg Vogel

**Affiliations:** Institute for Molecular Infection Biology (IMIB), University of Würzburg, 97080 Würzburg, Germany; Helmholtz Institute for RNA-based Infection Research (HIRI), Helmholtz Centre for Infection Research (HZI), 97080 Würzburg, Germany

## Abstract

Bacteriophages are the most abundant entities on earth and exhibit vast genetic and phenotypic diversity. Exploitation of this largely unexplored molecular space requires identification and functional characterisation of genes that act at the phage-host interface. To date, this is restricted to few model phage-host systems that are amenable to genetic manipulation. To overcome this limitation, we introduce a non-genetic mRNA targeting approach using exogenous delivery of programmable antisense oligomers (ASOs) to silence genes of both DNA and RNA phages. A systematic knockdown screen of core and accessory genes of the nucleus-forming jumbo phage ΦKZ, coupled to RNA-sequencing and microscopy analyses, reveals previously unrecognised proteins that are essential for phage replication and that, upon silencing, elicit distinct phenotypes at the level of the phage and host response. One of these factors is a ΦKZ-encoded nuclease that acts at a major decision point during infection, linking the formation of the protective phage nucleus to phage genome amplification. The non-genetic ASO-based gene silencing used here promises to be a versatile tool for molecular discovery in phage biology, will help elucidate defence and anti-defence mechanisms in non-model phage-host pairs, and offers potential for optimising phage therapy and biotechnological procedures.

## INTRODUCTION

Bacteriophages have become important study systems in molecular biology, biotechnology and medicine. This growing interest is driven by the richness of unexplored genes that emerge in the phage-host conflict (Bernheim & Sorek 2020, Mayo-Muñoz et al. 2024). Targeted mapping and characterisation of genes that act at the phage-host interface is key to understanding the major molecular players in a phage’s infection cycle and its ability to counter host defence, but the diversity and genetic intractability of many phages and their hosts creates a major challenge. While early work involved temperature-sensitive or other conditional mutants (Pires et al. 2016, Ofir & Sorek 2018, Mahler et al. 2022), chemical mutagenesis (Robins et al. 2013) and small RNAs (Sturino & Klaenhammer 2002), recent studies increasingly used CRISPR-Cas technology to inhibit phage gene expression (McDonnell et al. 2018, Piya et al. 2023, Adler et al. 2023, Sprenger et al. 2024). Notwithstanding the success of these techniques, a main barrier remains: Most of these approaches require genetic manipulation of the bacterial host. Yet, many phage hosts cannot be transformed or conjugated (Marsh et al. 2023) due to defence systems that target foreign DNA (Bernheim & Sorek 2022, Mayo-Muñoz et al. 2023). In addition, phages encode anti-CRISPR proteins or non-coding RNAs that can neutralise Cas enzymes (Bondy-Denomy et al. 2013, Camara-Wilpert et al. 2023). As a result, it remains difficult to map and characterise genes in the vast majority of phages, among them the intensely studied jumbo phage ΦKZ, which infects the major human pathogen, *Pseudomonas aeruginosa*.

ΦKZ-like jumbo phages are of great interest not only because of their potential for the treatment of recalcitrant *P. aeruginosa* infections via phage therapy (Chan et al. 2016, Cobián Güemes et al. 2023, Naknaen et al. 2024), but also because of their complex infection cycle that includes the sequential formation of membrane-and protein-bound compartments inside host cells. ΦKZ injects its genome together with a virion RNA-polymerase (vRNAP) inside an early phage infection (EPI) vesicle for immediate transcription of its genome (Armbruster et al. 2023, Antonova et al. 2024, Mozumdar et al. 2024). Subsequently, a subcellular structure referred to as the phage nucleus is formed, in which the non-viron RNAP (nvRNAP) continues transcription, and the phage genome is replicated and loaded into attaching phage capsids (Chaikeeratisak et al. 2017a, reviewed in Prichard & Pogliano 2024). These phage-induced cellular compartments shield the 280-kB dsDNA genome of ΦKZ from host defence mechanisms such as CRISPR-Cas and restriction enzymes (Malone et al. 2019, Mendoza et al. 2020) and make genetic gene silencing approaches challenging. Formation of the phage nucleus also necessitates mRNA export for cytosolic translation and import of *de novo* synthesised phage proteins.

Several conserved phage factors enable formation and organisation of the phage nucleus, e.g., the shell protein chimallin (ChmA) (Chaikeeratisak et al. 2017a, Laughlin et al. 2022, Nieweglowska et al. 2023), PicA, which mediates cargo trafficking across the phage nucleus (Morgan et al. 2024, Kokontis et al. 2024), and the tubulin-like protein PhuZ, which centres the phage nucleus in the middle of the cell and is involved in intracellular trafficking of newly assembled capsids (Erb et al. 2014, Chaikeeratisak et al. 2017a, Chaikeeratisak et al. 2017b, Chaikeeratisak et al. 2019). Nevertheless, major gaps remain in our understanding of the key decision points in the ΦKZ infection cycle. ΦKZ has ∼400 annotated protein-coding genes (Mesyanzhinov et al. 2002), but how many of them play an essential role in successful host take-over and the consecutive steps in the phage infection cycle remains unknown. This is largely due to a lack of genetic tools that permit targeted inhibition of ΦKZ genes.

In this study, we present a straightforward and broadly applicable non-genetic route to assess gene essentiality and function in phage-host interactions. Specifically, we employ programmable gene silencing at the RNA level via exogenous delivery of synthetic antisense oligomers (ASOs) into the bacterial cytosol. Such ASOs are typically 9-12 nucleobases in length and designed to sequester the ribosome binding site (RBS) or start codon (AUG) of a target mRNA to prevent synthesis of the encoded protein. They have been applied successfully in multiple bacterial species (Nielsen 2010, Vogel 2020, Pifer & Greenberg 2020). One popular ASO modality is peptide nucleic acid (PNA), which harbours a pseudo-peptide backbone that links the four natural nucleobases and protects the oligomer from nucleolytic and proteolytic degradation. For delivery across the bacterial envelope, the antisense PNA is fused to a cell-penetrating peptide (CPP, reviewed in El-Fateh et al. 2024) (**Fig. 1a**).

**Fig. 1.**
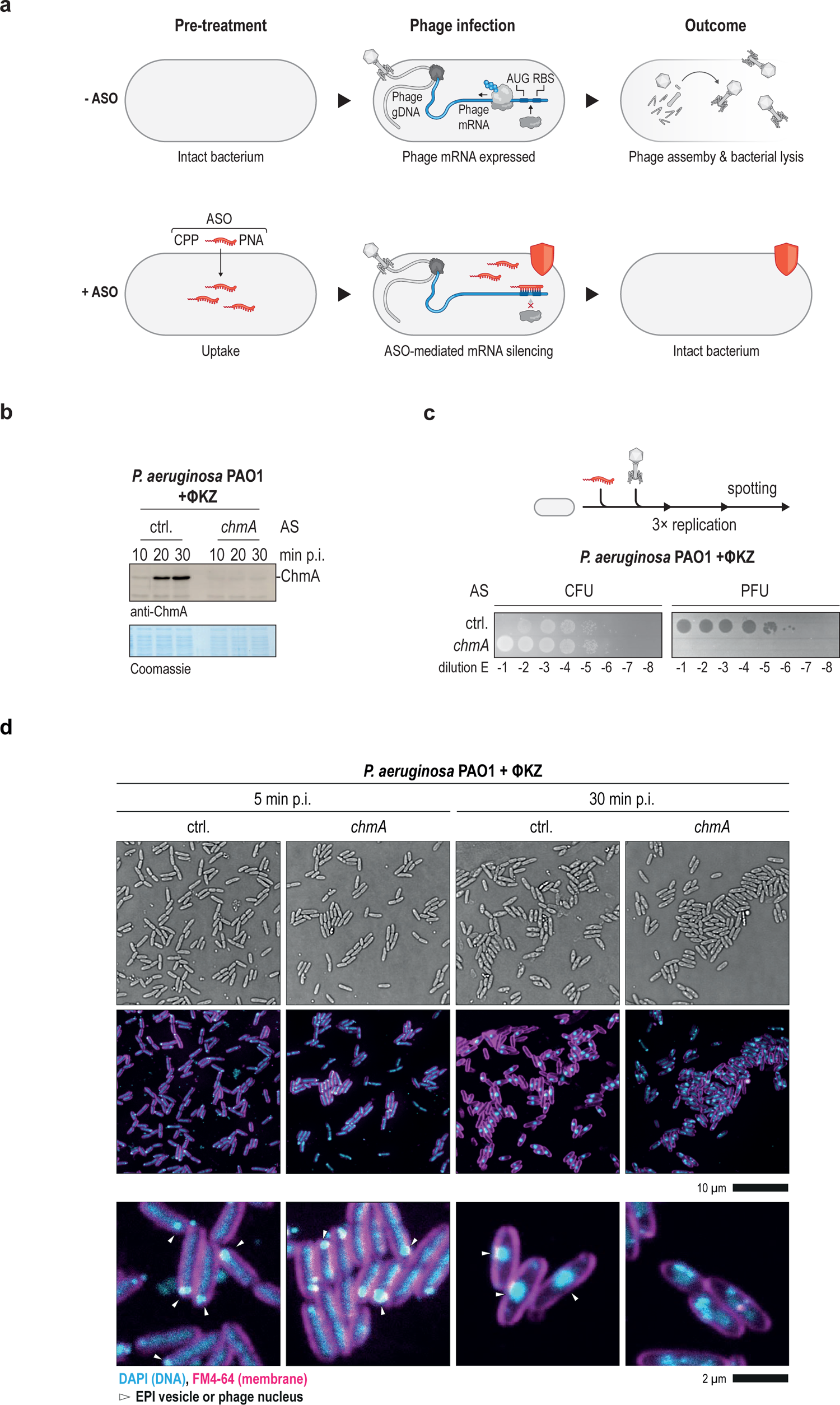
ASOs prevent ΦKZ phage replication in *Pseudomonas*. **a.** Antisense oligomers (ASO) are taken up by the bacterial cell mediated by cell penetrating peptides (CPPs). After phage infection, ASOs bind specific phage transcripts at the ribosome binding site (RBS) or the start codon (AUG) and prevent translation. If the protein encoded by the target phage mRNA is essential for phage propagation, no phage progeny is formed. PNA, peptide nucleic acid. **b.** PAO1 cells were pretreated with an ASO against *chmA* or a non-targeting control ASO (ctrl.). After ΦKZ infection with a MOI of 5, cells were harvested at the indicated time points and ChmA levels were determined by immunoblotting. AS: ASO to the indicated transcript. **c.** PAO1 cells were pretreated with an ASO against *chmA* or a non-targeting control. After ΦKZ infection at MOI 0.0001, cells were incubated to allow three rounds of replication, which leads to nearly complete lysis of cells. The resulting phage-cell suspension was spotted on LB plates to assess colony forming units (CFUs) and on LB plates with a lawn of PAO1 cells to assess plaque forming units (PFUs). A high number of phages leads to lysis of nearby bacterial cells after spotting, leading to complete lack of CFUs at low dilutions. **d.** PAO1 was treated with ASOs against *chmA* or a non-targeting control, infected with ΦKZ, followed by imaging. After chemical crosslinking, membranes are stained with FM4-64, DNA is stained with DAPI.

Here, we pioneer the application of ASO technology to score phage gene essentiality and to understand and break defence and counter-defence mechanisms in the phage-host interplay. Seeking to identify phage genes that serve key functions in the intricate infection process of phage ΦKZ in *P. aeruginosa*, we perform a systematic knockdown of ΦKZ genes coupled with phenotypic and transcriptome analyses. We discover numerous previously unknown essential genes with varying phenotypes, including a conserved phage ribonuclease that acts at a key decision point during progression from the early to the intermediate phase of the phage replication cycle. Our results suggest that more ΦKZ proteins than previously appreciated contribute to the subcellular organisation of the phage replication cycle.

## RESULTS

### ASOs can silence phage transcripts

Like other bacterial species, *P. aeruginosa* is amenable to ASO-induced gene silencing, as demonstrated by bactericidal ASOs that repress mRNAs encoding essential proteins of the transcription apparatus or in fatty acid synthesis (Howard et al. 2017, El-Fateh et al. 2024). Since phage mRNAs are translated outside the phage nucleus by cytosolic ribosomes, we hypothesised that they should be amenable to ASO-mediated silencing, too (**Fig. 1a**). Yet, it was unclear if ASOs could compete with the rapid overloading of the transcription-translation machinery by phage transcripts in infected host cells or whether mRNA synthesis inside the phage nucleus would shield these transcripts from ASO recognition. To establish a proof-of-concept, we designed ASOs against the mRNA of the essential chimallin protein ChmA (*ΦKZ054*), which is the main constituent of the phage nucleus (Chaikeeratisak et al. 2017a). As a control, we used an ASO that does not target any specific gene (‘non-targeting control’). Initially, we monitored ChmA protein levels after ASO treatment using a polyclonal antiserum raised against purified ChmA. We observed that 20 min post infection (p.i.), i.e., within the first round of replication, ASO knockdown fully prevented the synthesis of ChmA protein, even at a high multiplicity of infection (MOI) of 10 (**Fig. 1b**).

To test the functional effects of ChmA knockdown on phage replication, we assessed cell survival and phage progeny after multiple rounds of replication. This experimental set-up allowed us to amplify the effects on phage progeny, sample delayed replication times, and score effects that only become apparent in the next infection round. Initially, we optimised the experimental conditions. We adjusted the cell density (OD_600_ of 0.3), the ASO pre-treatment time (30 min), the concentration of ASOs (4-6 µM), the MOI (0.0001 to allow three rounds of replication until nearly complete host lysis), the time of spotting after infection (180 min; ∼ 3 replication rounds), and the length of the ASOs (10-11 nucleobases) (**Extended Data Fig. 1a-e**). Strikingly, knockdown of ChmA caused a strong reduction in phage progeny leading to a complete loss of plaques (**Fig. 1c**), demonstrating that sterilisation can be achieved with a *chmA*-targeting ASO.

Next, we sought to image phenotypic consequences of ChmA loss upon ASO knockdown. We therefore monitored progression through the phage replication cycle by imaging the formation of phage-derived cellular compartments. At 5 min p.i., the EPI vesicle was visible in both control and anti-*chmA* ASO treated cells (**Fig. 1d**), indicating successful infection. At 30 min p.i., the phage nucleus had formed in the control sample, evident as a central DNA-containing structure. By contrast, ASO-based knockdown of ChmA prevented the appearance of a recognizable phage nucleus. Instead, we observed smaller cellular structures that contained DNA, which we interpret as multiple EPI vesicles resulting from the high MOI of 10, necessary to achieve a synchronised infection. Hence, in the absence of ChmA, no phage nucleus forms, and this likely arrests the phage replication cycle (**Fig. 1d**). This agrees with observations made in *E. coli* infected with the phage Goslar after silencing of ChmA via CRISPR interference by antisense RNA targeting (CRISPRi-ART, Adler et al. 2023, Armbruster et al. 2023). The fact that ASO-based knockdown of ChmA completely prevented plaque formation (**Fig. 1c**) implies that the EPI vesicle is not able to support phage replication.

To extend the use of ASOs to study phage infection-related intracellular structures, we also silenced the *phuZ* mRNA encoding the phage spindle apparatus protein that positions the phage nucleus in the middle of the cell (Erb et al. 2014). Knockdown of *phuZ* resulted in decentralisation of the phage nucleus towards the cell pole (**Extended Data Fig. 2a,b**), effectively phenocopying *phuZ* gene deletion (Guan et al. 2022). These observations demonstrate that ASO-based silencing of phage genes is effective and can be coupled to imaging of cellular structures that form throughout the phage replication cycle to study phage biology.

### ASO applications for the study of the phage-host interplay

To prove the versatility of ASOs in the study of phage-host interactions, we extended the approach to genetically intractable strains, targeting either phage or host genes. *P. aeruginosa* strain PaLo44 is a clinical isolate (NCBI, PRJNA731114) that like many other strains isolated from patient samples has been refractory to transformation using classical approaches of bacterial genetics, such as electroporation (**Extended Data Fig. 2c**). Therefore, these strains are not amenable to CRISPR-based approaches. Using ASOs to target *chmA,* we were able to inhibit ΦKZ replication in PaLo44 with the same efficiency as observed in the *P. aeruginosa* lab strain PAO1 used above (**Fig. 2a**). Importantly, this clinical isolate—unlike PAO1—is not susceptible to infection with a ΦKZ phage lacking the *ΦKZ014* gene, which encodes a ribosome-associated protein that appears to circumvent a PaLo44-specific defence system (Gerovac et al. 2024). Using ASO-based knockdown of *ΦKZ014* (**Fig. 2b**), we were able to render PaLo44 resistant to ΦKZ as previously observed with an engineered Δ*ΦKZ014* phage (Gerovac et al. 2024). These data suggest that ASOs can be used to screen the impact of phage factors in divergent hosts without the need for genetic engineering.

**Fig. 2.**
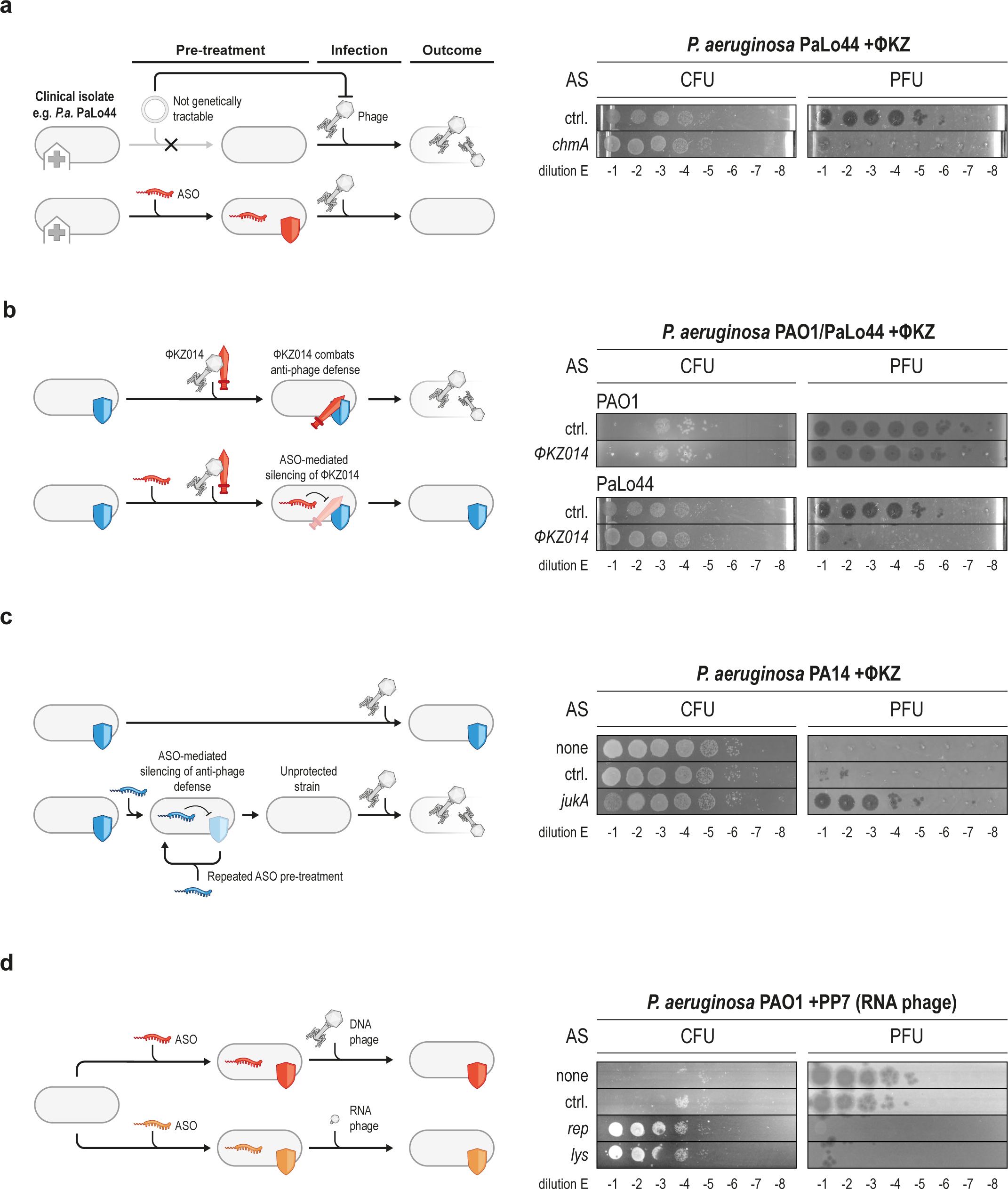
ASO applications for the study of the phage-host interplay. (Left) Schematic representation of potential ASO applications, such as targeting clinical isolates that are often genetically intractable **(a)**, protecting bacteria from phage infection by silencing phage genes that overcome an anti-phage defence system **(b)**, sensitising bacteria to phage infection by silencing a bacterial anti-phage defence system **(c)**, targeting RNA phages **(d)**. Anti-phage defence systems are depicted as shields, phage proteins that overcome bacterial defence as swords (Right) **a.** PaLo44 was treated with an ASO against *chmA* or a non-targeting control, cells were infected with ΦKZ at a MOI of 0.0001 and CFU/PFU were determined after 3 h incubation. **b.** PAO1 and PaLo44 cells were treated with ASOs against the phage transcript *ΦKZ014* followed by ΦKZ infection and CFU/PFU determination. **c.** PA14 cells were treated consecutively with an ASO against *jukA* followed by infection with ΦKZ and CFU/PFU determination. **d.** PAO1 cells were treated with ASOs against replication (*rep*) and lysis (*lys*) genes of the RNA phage PP7 followed by infection with PP7 and CFU/PFU determination. ctrl., non-targeting control.

Host-encoded defence systems are a major barrier to phage replication, but their genomic disruption is challenging. ASOs could be an alternative approach to inhibit their expression. To test this, we selected the JukAB defence system in the *P. aeruginosa* strain PA14, which rescues the cell by preventing early phage transcription, DNA replication, and nucleus assembly (Li et al. 2022). Since the *jukAB* locus is constitutively active, we inhibited the production of *jukA* prior to phage infection via repeated addition of a complementary ASO. ASO-mediated silencing of the *jukA* mRNA rendered *P. aeruginosa* PA14 bacteria susceptible to ΦKZ and allowed phage replication (**Fig. 2c**), indicating successful knockdown of the defence system. These results demonstrate that ASOs can be used to inhibit phage defence systems to study their composition and sensitivity in their native host and to propagate phages in otherwise non-permissive (pathogenic) strains.

RNA phages are an understudied class of bacterial viruses (Callanan et al. 2018), although knowledge of their diversity is expanding owing to novel techniques that reduce the sampling bias that normally favours discovery of DNA phages (Neri et al. 2022). To test the applicability of ASO technology to RNA phages, we selected the model RNA phage PP7, which also infects *P. aeruginosa* PAO1. We targeted the replication (*rep*, RNA-dependent RNA polymerase, RdRP) and lysis (*lys*) genes of PP7 with ASOs and completely abrogated plaque formation (**Fig. 2d**). Thus, our ASO-based knockdown approach is applicable to both DNA and RNA phages.

Overall, these experiments highlight that ASO-based gene silencing is a powerful technique to study many aspects of phage-host interactions, including in clinical isolates and strains that are not amenable to genetic manipulation. The technique can be applied to both host and phage genes and is agnostic to a phage’s type of genome. Moreover, we predict ASO-mediated silencing could be easily integrated into existing phage engineering methods and serve as a platform technology to modify phages or sensitise pathogens for phage therapy.

### Systematic ASO screen for factors important for phage replication

After establishing proof-of-concept, we reasoned that the programmable feature of ASOs would allow us to rapidly scale up this approach to screen for genes critical for phage replication. ΦKZ features a large genome with 377 annotated genes (NCBI), of which 85% have no predicted function (Mesyanzhinov et al. 2002, De Smet et al. 2017). This paucity of functional knowledge extends also to the much smaller core genome of *Chimalliviridae*, which consists of seven blocks of genes with likely interlinked functions, plus five independent genes (Prichard et al. 2023). To identify genes of ΦKZ that are essential for its replication, we screened 75 core and annotated genes (omitting most of the structural genes) using ASO-based knockdown and CFU/PFU readout in the model host strain *P. aeruginosa* PAO1.

Overall, ASO-mediated knockdown of one third of these genes (24) led to a strong effect (++, +++) on phage replication in CFU/PFU assays with multiple log of PFU reduction and CFU recovery. As expected, several ASOs that target factors known to be essential for phage replication, such as *chmA* (*ΦKZ054*), the nvRNAP subunits (*ΦKZ055* and *-068*) and the major head protein (*ΦKZ120*) triggered very strong effects (+++) (**Fig. 3** and **Extended Data 3a,b, Supplementary Data 1**). Silencing of PicA *(ΦKZ069*), which is required for protein import into the phage nucleus (Morgan et al. 2024, Kokontis et al. 2024) abrogated plaque formation as well. We also observed strong effects upon knockdown of uncharacterised genes. For example, silencing of *ΦKZ049*, a gene that encodes a SH3 domain protein suggested to be associated with the phage DNA polymerase (Iyer et al. 2021) reduced plaque formation substantially (**Extended Data 3b**). Similarly, knockdown of *ΦKZ042,* which encodes a conserved but uncharacterised protein, the hypothetical proteins *ΦKZ174/-177*, and the early expressed transcript *ΦKZ186* all strongly reduced plaque counts (**Extended Data 3b**). Knockdown of other genes produced milder effects, i.e., reducing plaque formation only 2-3 orders of magnitude, but nevertheless showing host protection from lysis as judged by CFU counts. This phenotype was observed for *ΦKZ144*, the gene encoding endolysin, which is vital for the final release of phage progeny from the cell (Briers et al. 2007, **Extended Data 3b**).

**Fig. 3.**
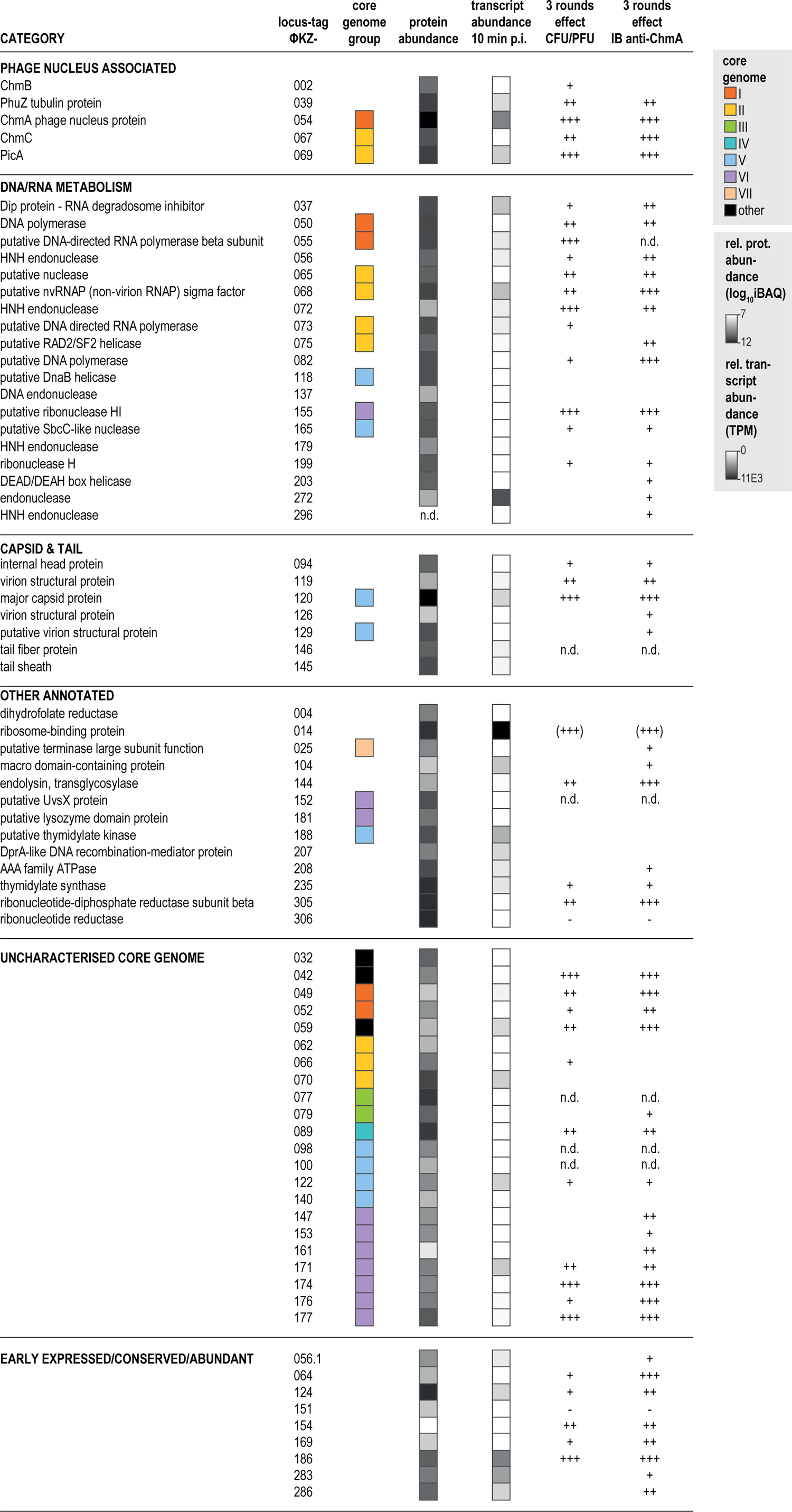
Screen for essential phage genes reveals factors that are important for ΦKZ replication. The impact of phage gene knockdown upon ASO treatment was evaluated by CFU/PFU determination and immunoblot detection of ChmA protein after three rounds of replication. The association of targets to core genome blocks in *Chimalliviridae* (Prichard et al. 2023), transcript abundance at 10 min p.i. and protein abundance (average values at 10, 20, 30 min p.i.) based on (Gerovac et al. 2024) are also shown. The PFU/CFU effect of each respective knockdown is depicted as no effect (), weak (+), effective (++), very effective plaque reduction (+++) and increased plaque levels (-); for details see **Extended Data** Fig. 3a. A minimum two ASOs were tested per gene, the stronger effect is shown. Toxic ASOs were omitted. n.d. not determined, e.g. if both ASOs were toxic. ΦKZ014 knockdown caused an abrogation of plaques only in the PaLo44 strain and is given in brackets.

In addition to CFU/PFU quantitation, we used ChmA levels as an additional readout, quantifying ChmA protein by immunoblotting after three rounds of replication. We observed reduced levels of ChmA for an additional 11 genes, although we saw no substantial changes of the infection rate as measured via CFU/PFU (**Fig. 3**). Examples include a putative RAD2/SF2 helicase (*ΦKZ075*), a predicted DEAD/DEAH box helicase (*ΦKZ203*), the macro domain-containing protein *ΦKZ104*, as well as the uncharacterised core genes *ΦKZ147, -153, -161, -176*, and *-161*, and the non-core genes *ΦKZ056.1, -283*, and *-286*. These genes can be expected to benefit phage fitness in more competitive situations such as non-laboratory environments or in *P. aeruginosa* strains with a different repertoire of defence systems.

We also observed indirect and potential off-target effects upon ASO treatment. For example, some ASOs were toxic to *P. aeruginosa* (22/176 of all tested ASOs, **Extended Data 3a,c**). Other ASOs increased PFUs by one order of magnitude, indicating that the targeted proteins, such as ΦKZ151 and -306, are negative regulators of phage replication or that possible off-target effects on the host may be beneficial for phage replication (**Extended Data 3c**). We also observed 4 ASOs that caused phage plaques without phage infection. This is indicative of activation of a prophage, most likely the integrative filamentous phage Pf4 (Knezevic et al. 2015, Gavric & Knezevic 2022, **Extended Data 3c**).

In summary, our screening approach yielded 45 phage proteins whose knockdown caused varying effects on phage propagation, and which offer promising leads for in-depth studies using ASO treatment in combination with phenotypic or multi-omics readouts.

### RNA-seq after ASO knockdown reveals molecular phenotypes of phage genes

Phage infection is a fine-tuned process that affects different cellular pathways, but not all will result in a macroscopic phenotype. We reasoned that more sensitive, global methods such as RNA-seq could reveal ’molecular phenotypes’ after ASO-mediated phage gene knockdown, defined as specific transcriptional dysregulation in infected cells (Westermann et al. 2016, Putzeys et al. 2024).

To establish this approach for ΦKZ, we first performed high-resolution RNA-seq after ChmA knockdown, which arrests the phage replication cycle at the level of the EPI vesicle (**Fig. 1d**, Armbruster et al. 2023). In seeking to analyse both the phage and host transcriptomes, total RNA was extracted from bacteria at 10, 15, 20, 25, 35 min after ΦKZ-infection. In a principal component analysis (PCA) of the RNA-seq data, the trajectory of the data from non-targeting ASO control samples revealed the transcriptional response throughout the phage replication cycle (**Fig. 4a, Supplementary Data 2**). Under these conditions, 10 min p.i. phage transcripts represented ∼45% of all sequenced reads that map to coding sequences and this increased to ∼70% at 35 min p.i. (**Extended Data Fig. 4a,b**, in agreement with Gerovac et al. 2024, Danilova et al. 2020, Ceyssens et al. 2017). However, upon *chmA* knockdown, reads of phage genes expressed at intermediate times were strongly depleted from 15 min p.i. onward, indicating that formation of the phage nucleus is required for expression of these genes by the nvRNAP (**Fig. 4b, Extended Data Fig. 4b-d,** classes E, F). Reassuringly, phage-encoded tRNAs were not detected after *chmA* knockdown either (**Extended Data Fig. 4c**), which is consistent with their dependence on the nvRNAP. Several other transcripts showed prolonged expression after *chmA* knockdown, indicating potential feedback inhibition from the phage nucleus (**Extended Data Fig. 4d**, classes A, B). Interestingly, the *chmA* transcript itself was unaffected by the ASO treatment, suggesting that in this case the successful translational inhibition does not entail mRNA depletion, as previously observed for several bacterial mRNAs targeted with ASOs in *Escherichia coli* (**Fig. 4b**, Popella et al. 2022).

**Fig. 4.**
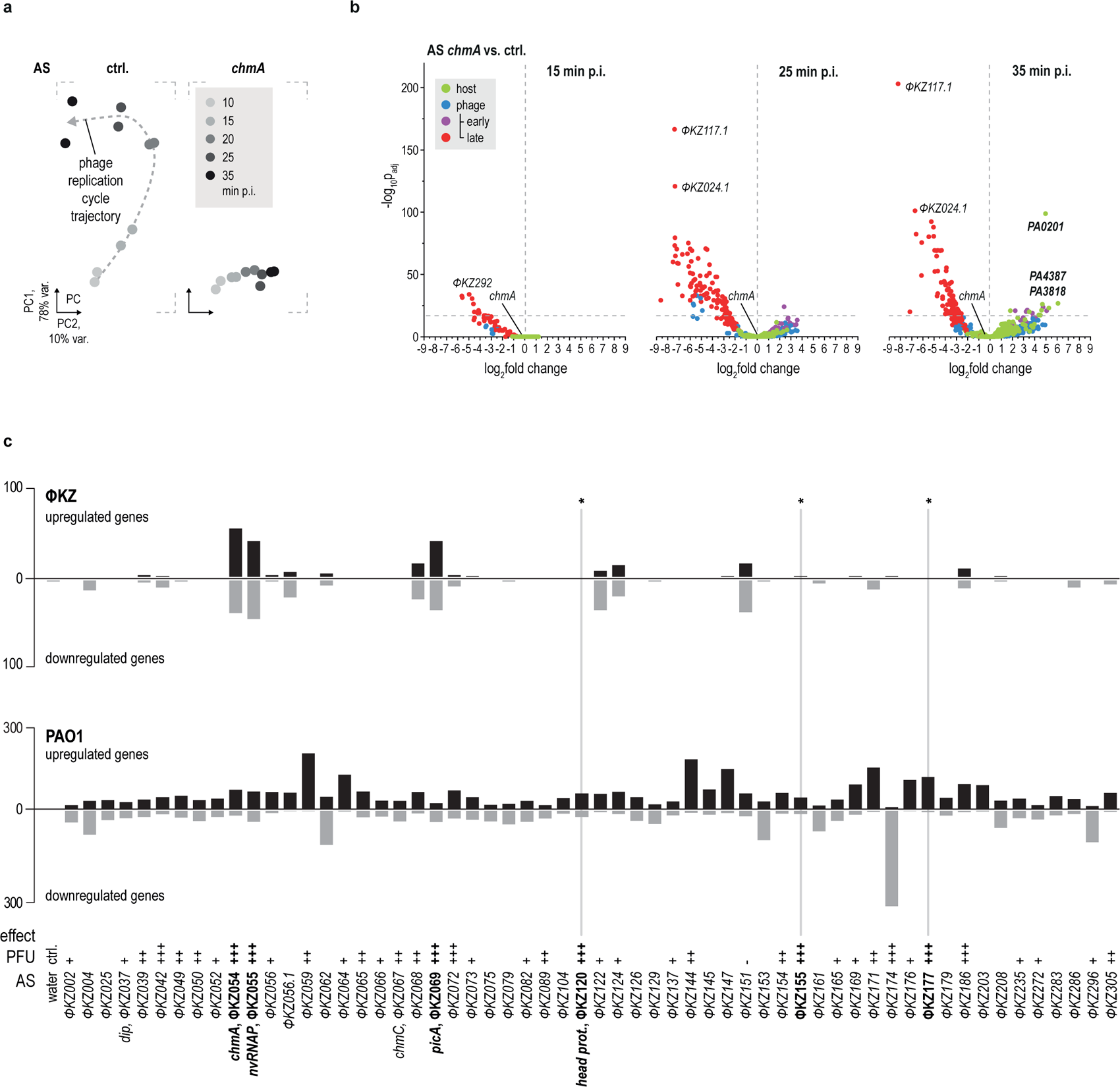
Transcriptional response after knockdown of ΦKZ core genes. **a.** ChmA was inhibited in ΦKZ-infected PAO1 cells and the transcriptome was sequenced at the indicated time points. To reduce the dimensionality of the dataset, we projected the dataset based on transcript abundance of each annotated gene on two dimensions via principal component (PC) analysis. Each dot represents a replicate, time points are shaded in grey ctrl., non-targeting control ASO. **b.** Transcript enrichment between ASO-knockdown vs non-targeting control ASO (ctrl.) is depicted in a volcano plot at indicated time points. Duplicates were merged by geometrical averaging and P-values were calculated by the Wald test using DESeq2. **c.** PAO1 cells were treated with ASOs against the indicated phage transcripts, infected with ΦKZ at a MOI of 5 and the transcriptome was analysed by RNA-sequencing at 30 min p.i.. Bars indicate the counts of ΦKZ (top) or PAO1 genes (bottom) that were affected in abundance (Log2FC>2, <-2, >100 reads in sum over all knockdown experiments at 30 min p.i.). For each target gene one replicate was sequenced. The PFU/CFU effect of each respective knockdown is depicted as no effect (), weak (+), effective (++), very effective plaque reduction (+++) and increased plaque levels (-), as in Fig. 3 and **Extended Data Fig. 3**. A minimum two ASOs were tested, the stronger effect is shown. Toxic ASOs were omitted. * Indicates the three genes whose knockdown led to a strong reduction in plaque formation but had no effect on the phage transcriptome.

On the host side, under control conditions, we did not observe a substantial transcriptional response at later stages of infection compared to 10 min p.i., when host-takeover is considered to be completed (**Extended Data Fig. 4b**). To test if this lack of a late host response is facilitated by the phage nucleus, we investigated host transcriptional changes upon *chmA* mRNA silencing. The absence of ChmA resulted in upregulation of 13 host transcripts out of ∼3,637 detected transcripts in *P. aeruginosa* at 35 min p.i.. Among these genes were *PA0201,* which encodes a hypothetical hydrolase; the membrane protein FxsA, which is linked to phage exclusion and may prevent superinfection (Cheng et al. 2004, Bondy-Denomy et al. 2016) and the type III secretion system regulator SuhB, which is important for *Pseudomonas* virulence (Li et al. 2013). Hence, even in the absence of a phage nucleus, the host response to the EPI vesicle is limited, which proves the protective nature of this structure.

Based on these data, we reasoned that transcriptomics is a sensitive readout that should allow the discovery of potential functional links between phage factors based on correlated transcriptomic profiles after their knockdown. We therefore extended the RNA-seq analysis to evaluating the consequences of ASO-mediated silencing of 58 genes from the screen above (**Fig. 4c, Supplementary Data 3**). At 15 min p.i., we observed diverse effects on phage transcript levels of genomic islands that did not cluster clearly (**Extended Data Fig. 5a**). At 30 min p.i., ASO-mediated depletion of ChmA, the nvRNAP subunit ΦKZ055, the nvRNAP sigma factor ΦKZ068, PicA (ΦKZ069), ΦKZ056.1, -122, -124, and -151 showed a strong effect on phage transcription (**Fig. 4c**). Of these, ChmA, the nvRNAP and PicA stood out because they strongly clustered based on ΦKZ transcript level alterations (**Extended Data Fig. 5b**). The fact that knockdown of PicA caused the same transcriptional phenotype as knockdown of ChmA and nvRNAP suggests that the nuclear import of nvRNAP subunits might be dependent on the recently discovered PicA protein importer (Morgan et al. 2024, Kokontis et al. 2024).

On the host transcriptome, we observed diverse responses upon knockdown of phage proteins. For example, knockdown of ΦKZ174, which abrogated phage replication, was accompanied by a strong downregulation of ∼300 PAO1 transcripts, i.e., almost 5% of all detected *P. aeruginosa* mRNAs (**Fig. 4c, Extended Data Fig. 5c-e**). While the function of the ΦKZ174 protein is unknown, we note that many of the genes repressed in its absence serve metabolic functions, suggesting that ΦKZ174 might act to prevent infected bacteria from metabolic shutdown. Equally interesting are the knockdowns of ΦKZ073 and -082, which have no general effect on the PAO1 transcriptome but specifically induce transcription of the Pf4 prophage locus (**Fig. 4c, Extended Data Fig. 5f**). Since awakening of the Pf4 filamentous bacteriophage can result in partial lysis of the bacterial population (Petrova et al. 2011), ΦKZ might use the ΦKZ073 and ΦKZ082 proteins as part of a specific inhibitory mechanism against the Pf4 prophage to ensure its own propagation. In summary, coupling ASO knockdown to RNA-seq revealed distinct transcriptional profiles of perturbed ΦKZ genes and allowed to cluster ΦKZ factors with different putative perturbation modes.

### ΦKZ155 is a phage nucleus-located RNase H essential for phage replication

The major goal of our ASO-based knockdown screen was to identify ΦKZ factors with key roles in the progression of the phage replication cycle, as defined by locking infection in a defined state. In this regard, we noticed a group of genes, i.e., ΦKZ120, ΦKZ155, and ΦKZ177, whose silencing strongly reduced plaque formation but hardly altered phage gene expression within the time course of our analysis (**Figs. 4c, 5a**). This can be rationalised for ΦKZ120 (major head protein) and ΦKZ177 (function unknown), which are expressed during later stages of infection (**Supplementary Data 3**). By contrast, ΦKZ155 is expressed at 20 min p.i. and its expression is dependent on ChmA (**Extended Data Fig. 5g**). This suggests that ΦKZ155 plays an important role after the initial formation of the phage nucleus.

**Fig. 5.**
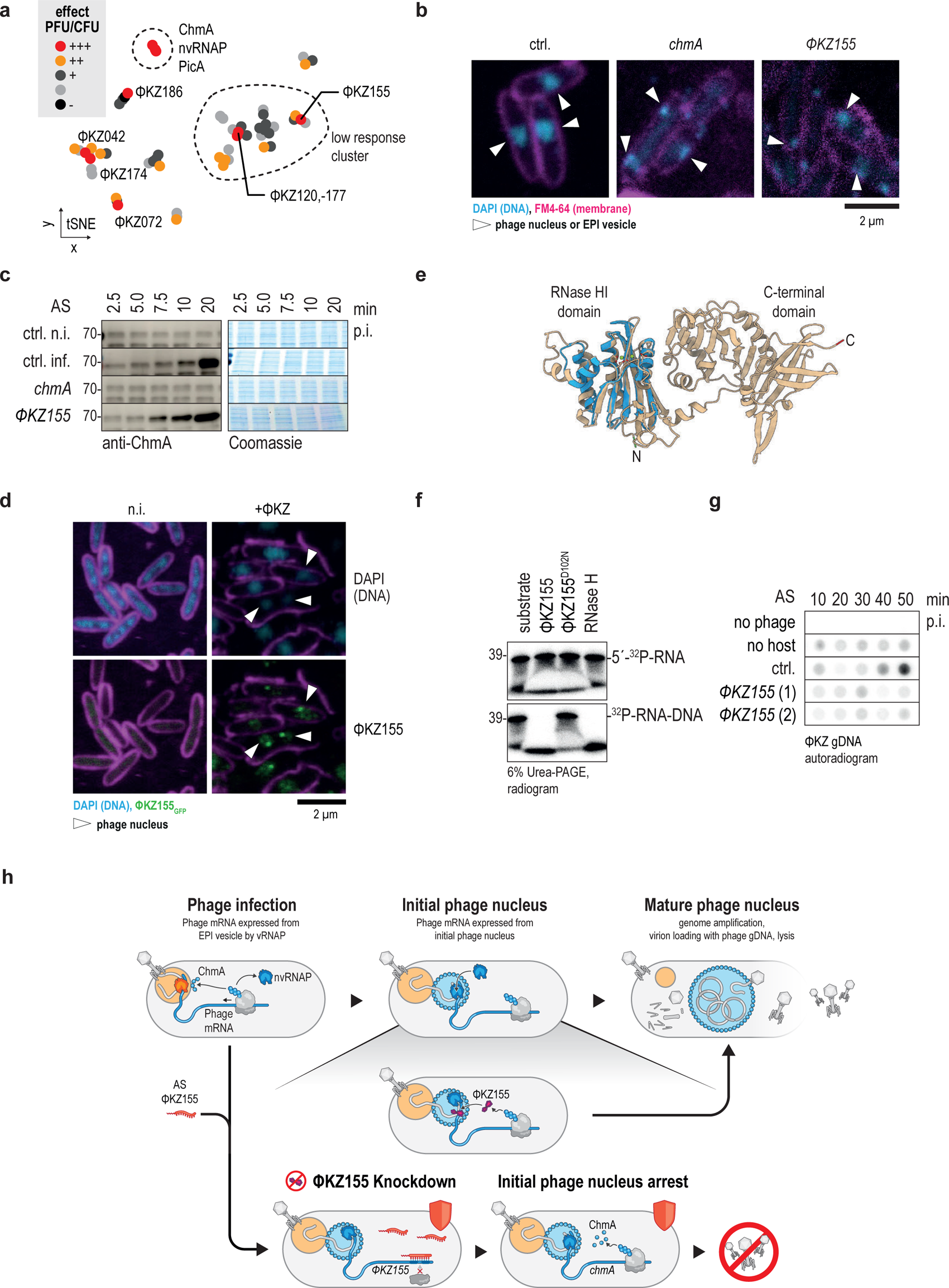
ΦKZ155 is important for phage nucleus maturation and phage genome amplification. **a.** Log2FC values for each ΦKZ gene from Fig. 4c and **Extended Data Fig. 5b** were clustered by tSNE. Clusters were indicated by a dashed line. **b.** PAO1 cells were treated with a non-targeting control ASO (ctrl.) or an ASO against ChmA or ΦKZ155 and infected with ΦKZ. Cells were chemically crosslinked after 35 min p.i. and stained with DAPI and FM4-64 to visualise DNA and membranes, respectively. **c.** PAO1 cells were treated like in (a) and harvested at indicated time points for SDS-PAGE and Western blotting against ChmA with a polyclonal anti-ChmA antibody serum. Coomassie stained gels served as loading control. **d.** ΦKZ155_GFP_ was ectopically expressed from a plasmid in PAO1, followed by infection with ΦKZ. Cells were chemically crosslinked, stained, and imaged. **e.** The ΦKZ155 structure was predicted using the AlphaFold 3 server. The N-terminal domain aligns well with a predicted model of an RNase HI domain (AF-A0A2A2IBB4-F1, blue). N-terminus and C-terminus are coloured in green and red, respectively, magnesium ions are represented as green balls. **f.** RNA was 5’-[^32^P]phosphorylated and optionally duplexed with complementary DNA (bottom). ΦKZ155 and the catalytically dead D102N mutant were produced by *in vitro* translation and added to the oligomer for the cleavage reaction. Subsequently, the oligomers were analysed on an Urea-PAGE gel and autoradiographed. **g.** PAO1 cells were treated as in (a) and the cell culture was harvested at indicated time points for DNA extraction. The samples were dot blotted, RNA was eliminated by alkaline treatment. The phage genomic DNA was detected with the radiolabelled oligo probe JVO-23213 (complementary to the *chmA* open reading frame) and autoradiography. **h.** Model of ΦKZ155 function. (Top) The ΦKZ155 protein is imported into the nucleus where it plays a role in phage nucleus maturation, which is likely linked to phage genome amplification. After genome amplification, the phage genome is loaded into virions at the phage nucleus. (Bottom) Upon knockdown of ΦKZ155, the initial phage nucleus remains immature, although cellular ChmA levels are increased. The phage genome is not amplified, and phage infection is halted.

To better understand the role of ΦKZ155 during phage infection, we performed fluorescence microscopy imaging of phage-infected cells. To our surprise, knockdown of ΦKZ155 inhibited the formation of a large phage nucleus centred within the cell at 35 min p.i. (**Fig. 5b**), despite the fact that phage nucleus-dependent transcription of phage genes was unaltered (**Fig. 5a, Extended Data Fig. 5b**). Importantly, this effect was not due to reduced levels of the phage-nucleus protein ChmA (**Fig. 5c**). In fact, western blot analysis showed ChmA levels to be slightly elevated early during infection upon ΦKZ155 knockdown. To probe the subcellular localization of the ΦKZ155 protein, we ectopically expressed GFP-tagged ΦKZ155 and found that the protein localises inside the phage nucleus (**Fig. 5d**). Curiously, the ΦKZ155_GFP_ protein was barely detectable in the absence of phage infection, indicating that it requires the phage nucleus for its own stability. Therefore, ΦKZ155 is an essential phage factor that facilitates the switch towards phage nucleus maturation.

ΦKZ155 is part of the core set of proteins encoded by the nucleus-forming bacteriophage family (Prichard et al. 2023). Its predicted structure harbours an N-terminal RNase HI domain (UniProt, Pfam domain PF00075) and a well-structured C-terminal domain of unknown function (local distance difference test >0.9, AlphaFold3 server, Abramson et al. 2024, **Fig. 5e**). RNase HI domains are typically found in enzymes that recognize RNA-DNA heteroduplexes and specifically cleave the RNA strand (Moelling et al. 2017). To validate this predicted nucleolytic mode of action, we synthesised the ΦKZ155 protein in an *in vitro* translation (IVT) reaction and added RNA and RNA-DNA substrates. Both the ΦKZ155 protein and *E. coli* RNase H, included as a positive control, triggered RNA degradation when offered the RNA-DNA duplex, while single-stranded RNA was not processed (**Fig. 5f**). Moreover, the DNA-RNA duplex dependent processing by ΦKZ155 was lost when the conserved RNase HI catalytic residue Asp102 was mutated to Asn. These data experimentally establish that ΦKZ155 has intrinsic RNase HI-like activity.

A main cellular function of RNases H-type nucleases is in preserving DNA integrity, protecting the replication machinery against the topological challenge posed by RNA–DNA hybrids, commonly referred to as R-loops (Moelling et al. 2017). However, these nucleases also degrade RNA primers that initiate genome replication in both prokaryotes and eukaryotes, which suggested that ΦKZ155 might facilitate ΦKZ genome amplification. To test this, we quantified phage DNA levels by dot blot analysis in cells treated with a control ASO or a ΦKZ155-targeting ASO following phage infection. In this assay, increased phage DNA is detected 40 min p.i. in *P. aeruginosa* PAO1 treated with the control ASO. By contrast, we observed no such phage genome amplification after ASO-mediated silencing of ΦKZ155 (**Fig. 5g**). Thus, the lack of ΦKZ155 arrests the phage infection cycle at a crucial stage shortly after the initial phage nucleus is made, by stalling amplification of the phage genome (**Fig. 5h**). We predict that such defined arrested states achieved by ASO-based knockdown of specific phage genes will be useful in dissecting the molecular steps that govern key decision points in the progressing phage cycle.

## DISCUSSION

The ASO-mediated gene silencing reported in this study is expected to be of broad use in phage biology because it greatly enhances our ability to assess gene functions in phage-host interactions. While applied here to phages of *P. aeruginosa*, the general strategy is readily adaptable to other phages and hosts, including ones that are genetically non-tractable. As shown here, ASOs can be used to study many aspects of phage-host interactions, beyond scoring the essentiality of single genes. We demonstrate successful silencing of genes in RNA and DNA phages (including a nucleus-forming jumbo phage), suppression of a host-encoded phage defence system, applicability to clinical isolates, and scalability to enable systematic screens for essential genes of a phage of interest. We have also shown that the coupling of ASO-mediated gene silencing to sensitive readouts such as RNA-seq greatly extends the information output from such screens. Our sampling of a fifth of the genes of the model jumbo phage ΦKZ clearly demonstrates that many genes that lack predictable physiological or molecular functions produce macroscopic or molecular phenotypes when silenced. Given that our screen was performed in one growth condition and a single host, we are likely underestimating the proportion of functional genes.

It is also important to bear in mind that the ASOs used here were designed to translationally silence mRNAs, with the goal to suppress protein synthesis. However, there is a growing appreciation of noncoding RNA functions in phage-host interactions (Altuvia et al. 2018, Sprenger et al. 2024), and the ASO approach used here should be easily applicable to inhibiting phage-related regulatory small RNAs. Likewise, ASOs could be important for exploring the expanding class of minimal RNA replicators such as viroids and viroid-like covalently closed circular (ccc) RNAs, some of which are predicted to replicate in environmental bacteria (Lee et al. 2023, Zheludev et al. 2024). Further, we foresee applications in phage therapy, e.g., in the optimisation of production strains or phage cocktails, or in industrial settings, e.g., preventing starter cultures from detrimental phage infections. Such applications should greatly benefit from a main distinguishing feature of ASO-based knockdowns, which is that they obviate the need for generating a genetically modified organism (GMO) in the process.

Our screening data represent rich resources of interesting phage genes for further in-depth study and a source for effective ASO targets in the family of nucleus-forming jumbo phages. Our follow-up analysis of the conserved ΦKZ0155 protein revealed an essential phage-encoded ribonuclease and suggests a role of this protein in the temporal coordination of phage nucleus maturation and genome amplification. Proteins with RNase HI domains typically enable host genome amplification by providing RNA primers for leading strand synthesis (Itoh & Tomizawa 1980) or by degrading RNA primers (Moelling et al. 2017). Although the molecular mechanisms by which most phages replicate their genomes are unknown, it has been assumed that phage-related DNA-RNA duplexes are resolved by host-encoded RNase H. For example, an RNase HI encoded by phage T4 is only essential for genome amplification in RNase HI-deficient hosts (Hollingsworth & Nossal 1991, Hobbs & Nossal 1996, Mueser et al. 2010). By contrast, our results suggest that ΦKZ has evolved to become independent of its host for RNase H activity, perhaps because the phage nucleus excludes *P. aeruginosa* RNase HI proteins. Instead, we hypothesise that through acquisition of the ΦKZ155 gene and through targeting the encoded protein to the phage nucleus, ΦKZ and other jumbo phages have monopolised an RNase H activity to time genome amplification with maturation of the cellular structure that will eventually assemble the components for new viral particles. It will be intriguing to assess the molecular function of the C-terminal domain of ΦKZ155 and identify molecular interaction partners of this nuclease to understand the temporal and spatial control of this crucial step in the phage replication cycle. The transient state shortly after the phage genome is translocated from the EPI vesicle into the phage nucleus is poorly understood molecularly and structurally but now addressable by arresting this state with ASOs against ΦKZ155.

It is intriguing to note that ΦKZ encodes at least ten nucleases (2.5% of all annotated genes). Their targets and catalytic mechanisms remain largely unknown, except for ΦKZ179, whose homolog in phage ΦPA3 has just been shown to help ΦPA3 attack the ΦKZ genome when both phages infect the same *Pseudomonas* cell (Birkholz et al. 2024). In agreement with a specialised role in inter-phage warfare, knockdown of the ΦKZ179 nuclease did not show a phenotype in our assay where ΦKZ was the only phage present (**Fig. 3**). However, our screen did identify at least one additional candidate, ΦKZ072, which encodes a predicted HNH nuclease, with a strong knockdown phenotype in the range of ChmA or ΦKZ155 (**Fig. 3**). Interestingly, ΦKZ072 is expressed early and does not require phage nucleus formation for its expression, which suggests a role in early host take-over (**Supplementary Data 2**). Other candidates with milder phenotypes under certain assay conditions were ΦKZ056, ΦKZ165 and ΦKZ199. With the ability to inhibit their expression now in hand, it will be intriguing to unravel how these essential and non-essential nucleases act at different stages in the finely orchestrated phage replication cycle.

## Extended Data

**Extended Data Fig. 1.**
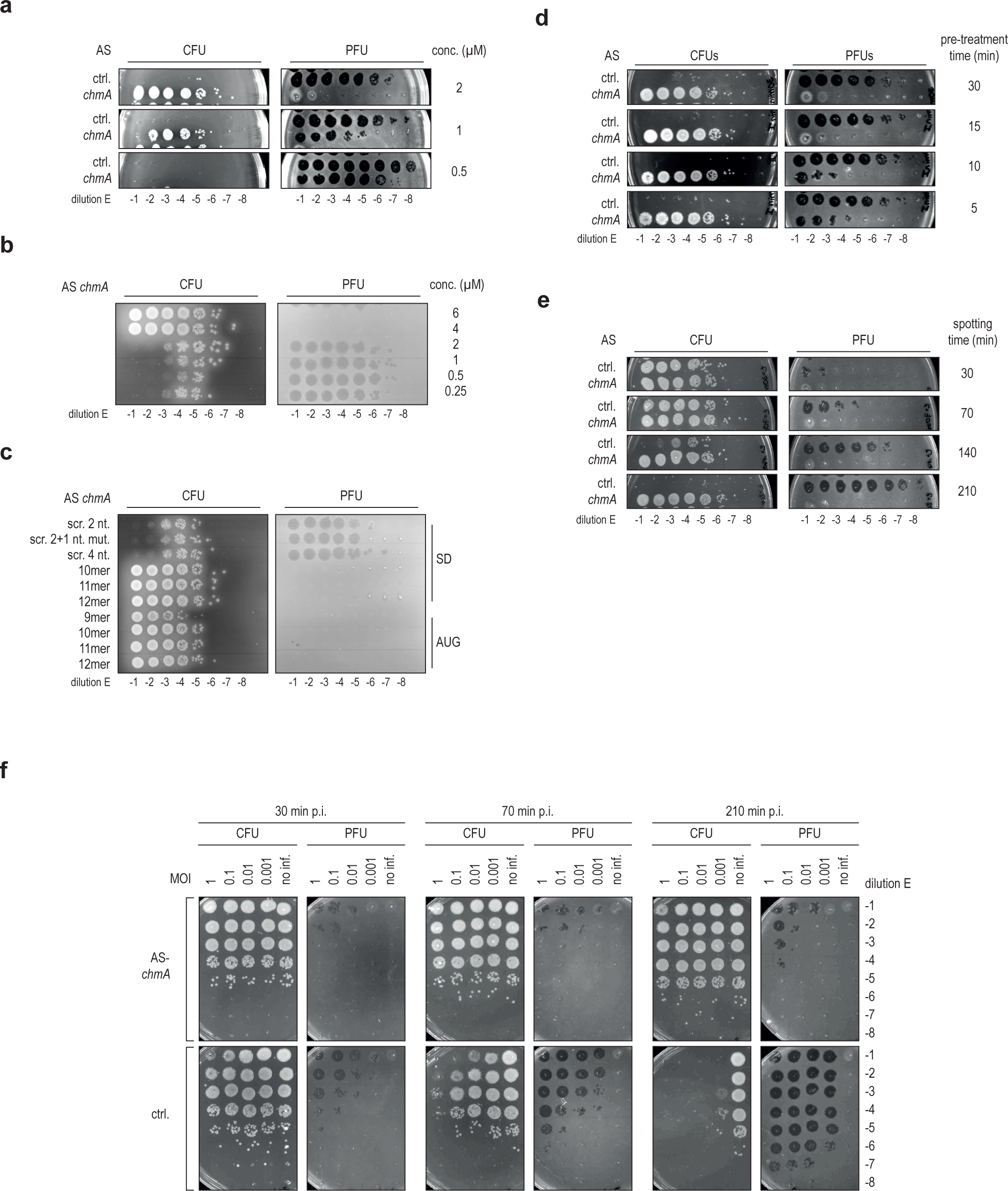
Optimization of ASO treatment. **a.** Effective ASO concentration was determined by pre-treating cells for 30 min with different ASO concentrations as indicated. Subsequently, cells were infected with ΦKZ and incubated for 30 min, which was followed by CFU/PFU readout. **b.** Effective ASO concentration was determined by pre-treating cells for 30 min with different ASO concentrations as indicated. Subsequently, cells were infected with ΦKZ and allowed to replicate for 3 h, which was followed by CFU/PFU readout. **c.** ASOs of different lengths and scrambled variants were tested. For the scrambled variants, two (2x) or four (4x) nucleotides were changed in their position, 3x represents 2x with one additional mutation; all changes were made in the central part of the ASO-sequence. ASOs were added at 6 µM. Subsequently, cells were infected with ΦKZ, allowed to replicate for 3 h, which was followed by CFU/PFU readout. SD Shine-Dalgarno sequence; AUG start codon. **d.** ASO pre-treatment time was tested at 5, 10, 15, and 30 min. **e.** Sampling time after infection was tested at below one replication round (30 min) and at one (70 min), two (140 min), and three (210 min) replication rounds, followed by CFU/PFU readout. **f.** MOI used for infection was tested at 1, 0.1, 0.01, and 0.001.

**Extended Data Fig. 2.**
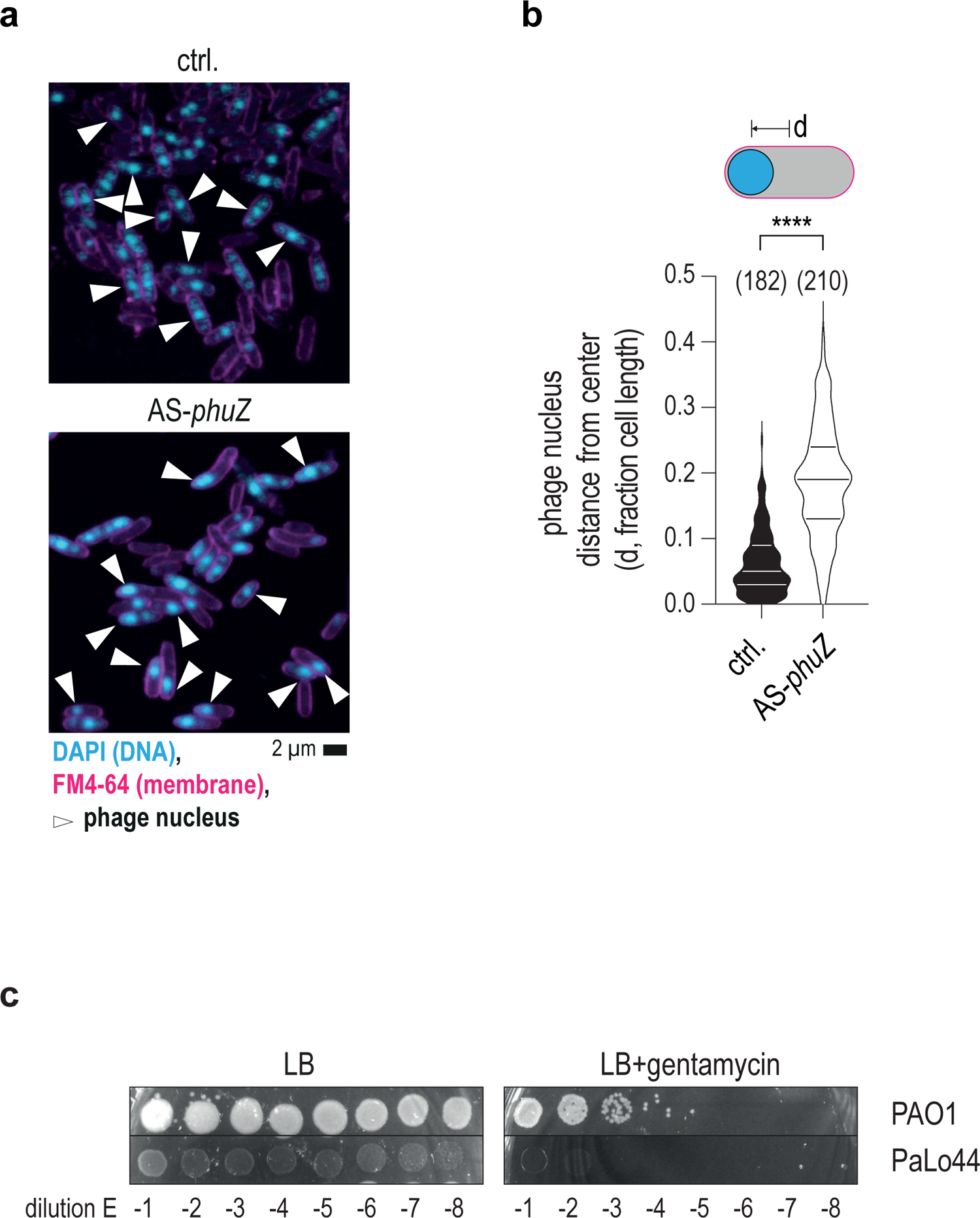
Knockdown of PhuZ decentralises the phage nucleus. **a.** Imaging of ΦKZ-infected PAO1 cells after ASO-based knockdown of the phage spindle apparatus gene *phuZ*. **b.** Phage nucleus distance from centre after *phuZ* knockdown. The data are based on one representative example of two independent experiments. The number of counted cells is indicated in brackets. **** p<0.0001, two-tailed Mann-Whitney test. **c.** Transformation efficiency of PAO1 and the clinical isolate PaLo44 with empty plasmid carrying a gentamicin resistance marker. After transformation, cells were spotted for CFU counting in a dilution series on LB plates with or without gentamicin.

**Extended Data Fig. 3.**
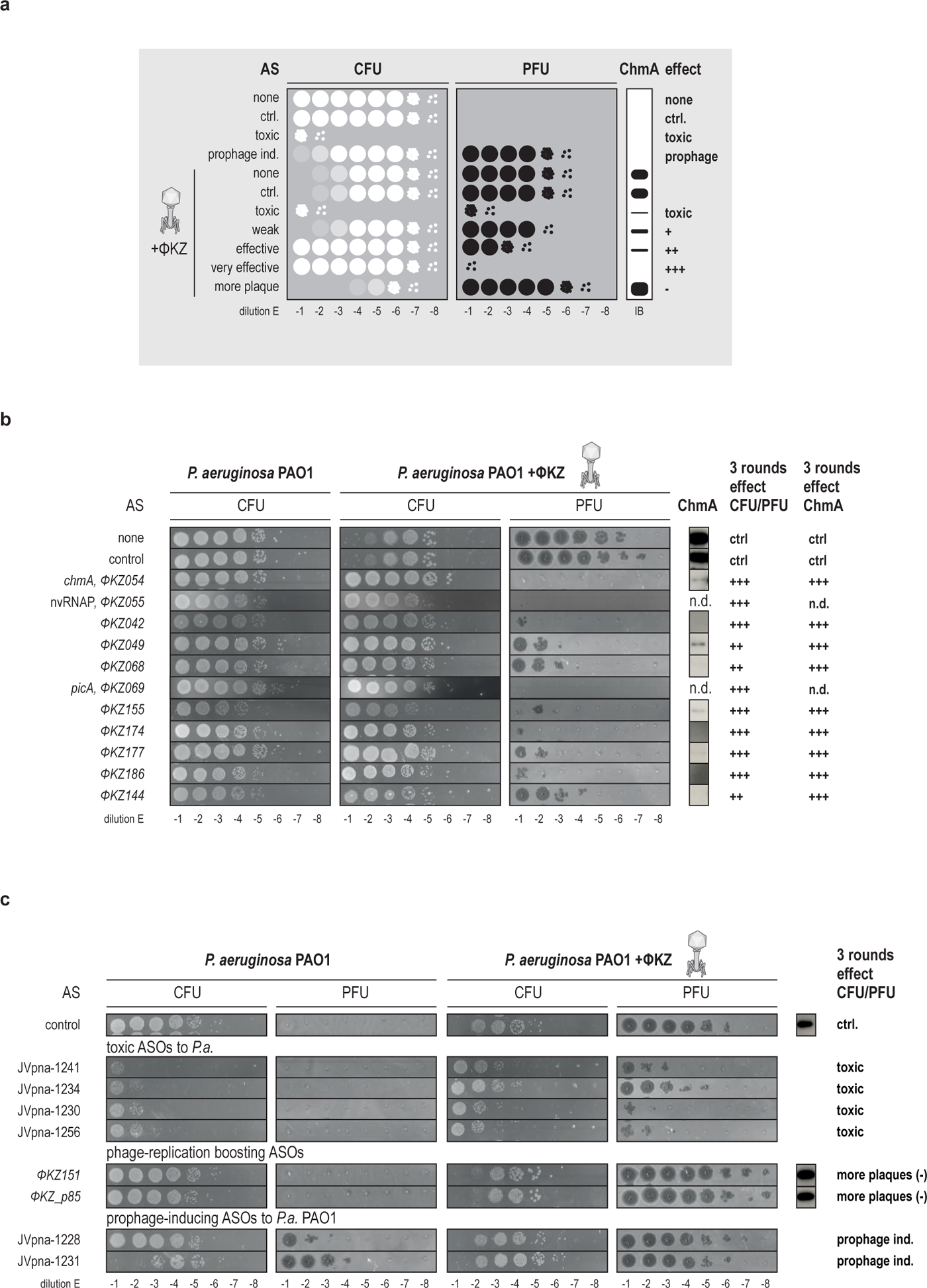
Potential off target effects caused by ASOs. **a.** Schematic representation of possible CFU/PFU readouts upon ASO treatment. ASOs can cause pleiotropic effects on CFU/PFUs and the level of ChmA at 30 min p.i.. Without phage infection, ASOs can be toxic to the host causing reduced CFU counts, or induce a prophage visible as PFUs and optionally reduced CFUs at low dilutions. Toxic ASOs will also cause reduced PFUs upon infection with ΦKZ. ASOs that showed one log reduction in PFUs and/or reduced ChmA levels were scored as weak (+). ASO that reduced PFUs by multiple logs, rescued CFUs in the first dilution, and/or depleted ChmA levels were scored as effective (++). ASOs that diminished PFUs down to the first dilution or completely abrogated PFUs, and/or depleted ChmA levels by more than 10-fold were scored as very effective (+++). Some ASOs caused increased plaque counts (-) and/or increased ChmA levels. **b.** CFU/PFU and ChmA levels upon knockdown of top candidates with a strong effect on phage replication. Scoring as described in (a). **c.** Examples of pleiotropic effects of ASO treatment. ASOs can be toxic for the host. These ASOs are denoted by their internal reference number because the effects might be unspecific and not related to the intended target gene. ASO treatment can result in more phage plaques (-). ASOs can induce prophages (prophage).

**Extended Data Fig. 4.**
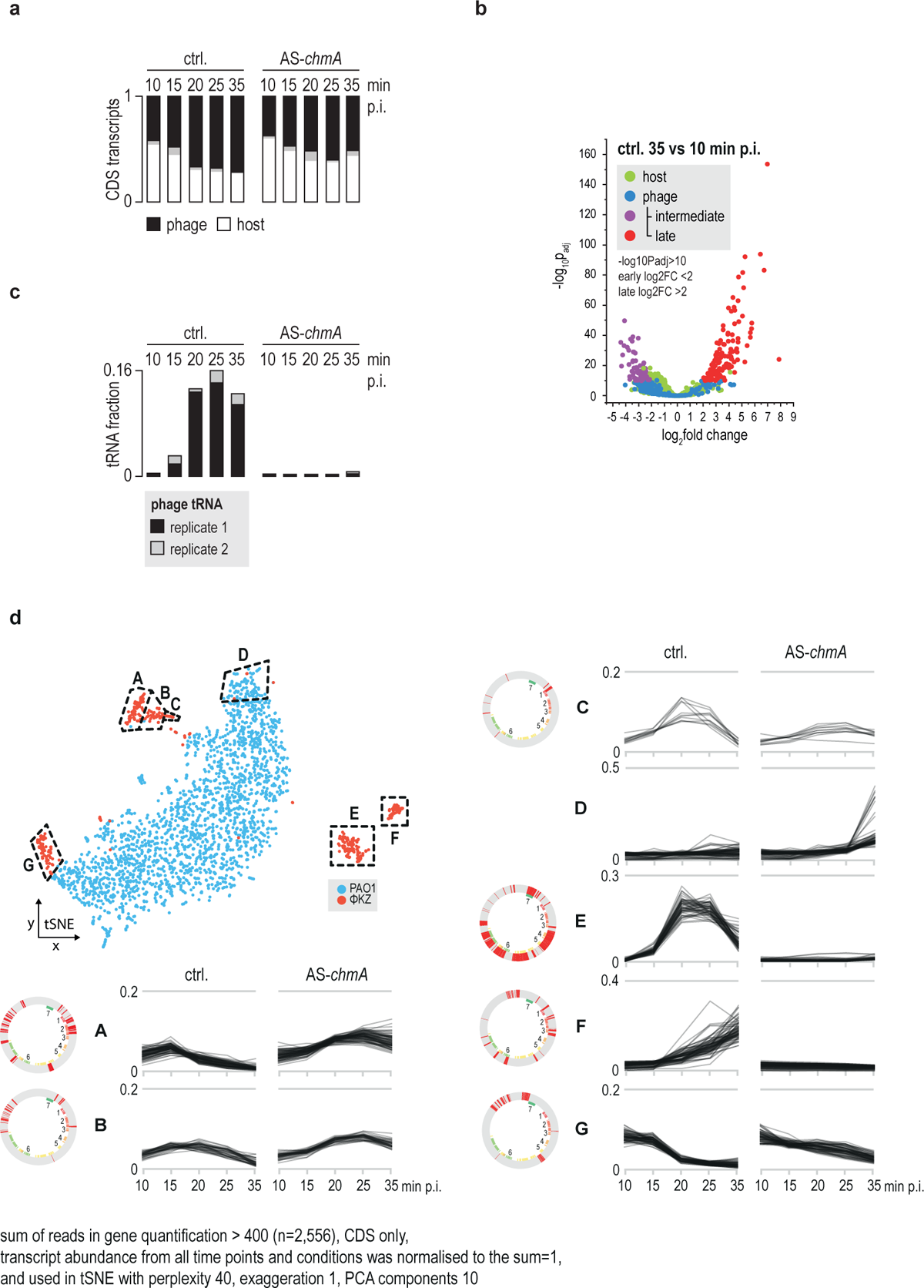
Early-intermediate gene classification and effects upon inhibition of ChmA. **a.** Relative quantification of transcripts that encode coding sequences (CDS) at indicated time points post infection after treatment with a non-targeting control ASO (ctrl) or a *chmA*-ASO. **b.** Enrichment of transcripts between 35 and 10 min post infection in the control samples. Phage genes were classified as intermediate (purple) upon depletion, and as late (red) upon enrichment. Other phage transcripts (blue) represent genes expressed early from the EPI vesicle or others, if the levels remained constant. Duplicates were merged by geometrical averaging and the P-values were calculated by the Wald test. **c.** Relative read counts for phage tRNAs in the tRNA fraction in the non-targeting control and ChmA knockdown condition; results are depicted in overlayed duplicates. **d.** Transcript levels of control and *chmA*-ASO experiments were normalised over the course of infection and clustered by tSNE. Individual clusters are represented together with genomic locations, core genes and blocks.

**Extended Data Fig. 5.**
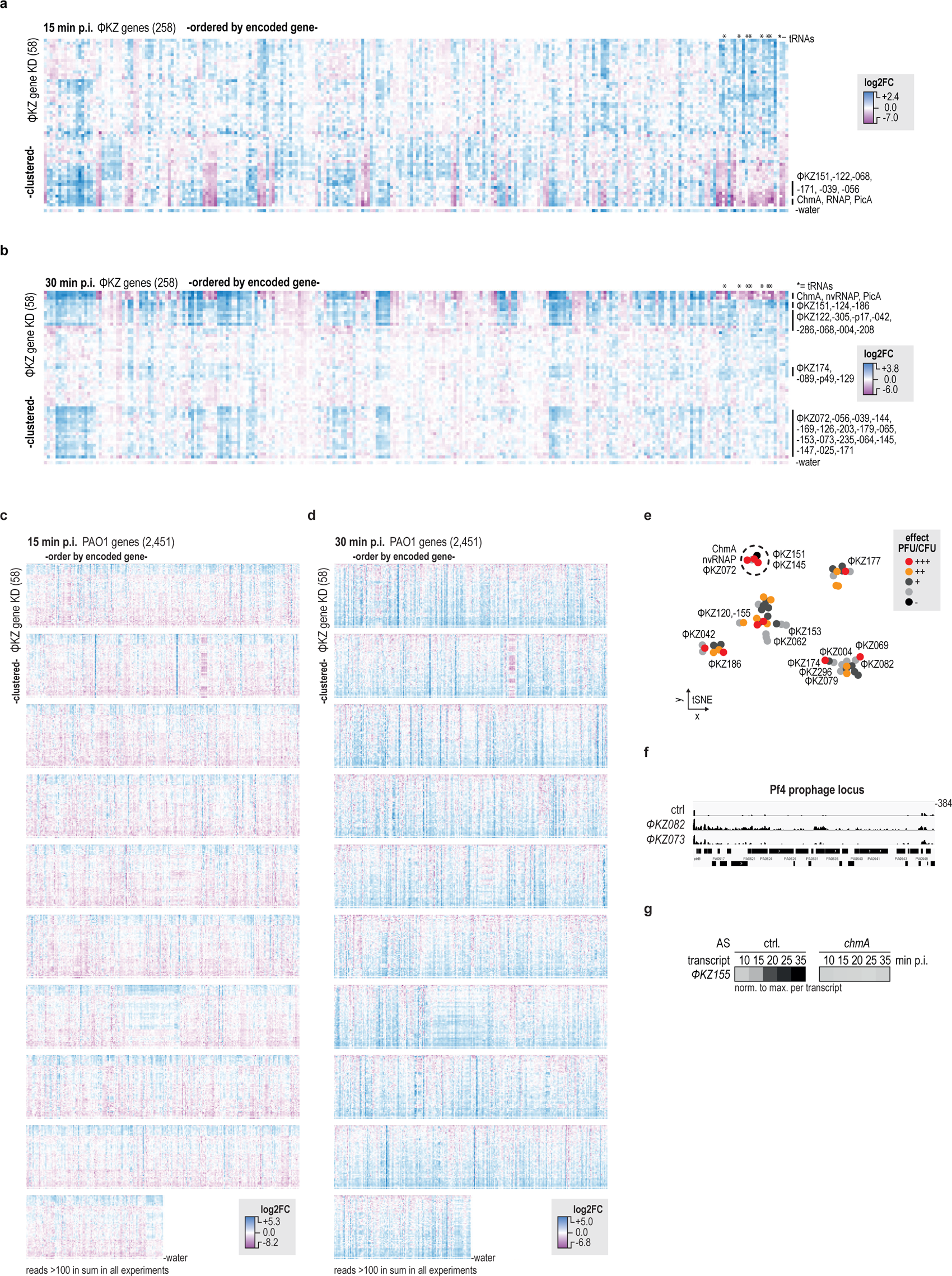
ASO-treatment reveals molecular phenotypes in host and phage transcript levels. **a., b.** Heatmap for Log2FC for transcripts from ΦKZ, 15 and 30 min p.i., respectively. **c., d.** Heatmap for Log2FC for transcripts from PAO1 at 15 min p.i. and 30 min p.i., respectively. **e.** tSNE clustering of Log2FC PAO1 transcript levels upon knockdown of indicated ΦKZ genes. Phage genes upon knockdown led to very effective plaque reduction (+++) and phage genes that showed strong phenotype in the host transcriptome (in Fig. 4c) are labelled. **f.** Read coverage of the Pf4 prophage locus in PAO1 upon knockdown of ΦKZ082 and -073. **g.** Transcript levels of *ΦKZ155* in non-targeting ctrl. and ChmA knockdown conditions (analysis based on data described in Fig. 4a**,b**).

## Methods

### ASOs

ASOs were designed against RBS and AUG regions using the MASON algorithm (Jung et al. 2023, https://mason.helmholtz-hiri.de) and the NCBI sequence and annotation files (ΦKZ: NC_004629.1, PP7: NC_001628.1). ASO length was set to 11-mers and the allowed mismatches for off-targets were set to 4. ASOs were selected based on the following scoring values: melting temperature (45-55°C), low purine percentage (25-35%) and few predicted off-targets in distinct translation initiation regions of the phage (<3). The scoring values for the used ASOs are given in **Supplementary Data 1**. At least two ASOs were designed for each targeted gene. The control ASO-sequence was GACATAATTGT (ctrl., JVPNA-79). ASOs were commercially ordered at Peps 4LS (Heidelberg) with a peptide-backbone (PNA) and a 5’-RXR (RXRRXRRXRRXRXB) CPP. The initial concentration was adjusted to 1 mM in water based on the specific extinction coefficient using absorption. ASOs were stored at -20 °C. Prior usage, ASOs were thawed at room temperature, then heated for 5 min at 50 °C and then cooled down at room-temperature.

### CFU/PFU assay

PAO1 (JVS-11761, DSMZ: DSM22644), PaLo44 (R. Lavigne lab, KU Leuven, Belgium), PA14 (R. Lavigne lab, KU Leuven, Belgium) were grown in LB media overnight at 37 °C and 220 rpm. Cells were inoculated 1:100 and grown in MH media at 37 °C and 220 rpm to an OD of 0.3 (absorption given at 600 nm). ASOs were added at 6 µM final concentration to 50 µl cultures and incubated for 30 min. Cells were infected with ΦKZ at an MOI 0.0001 and the cells were incubated for 3 h. 5 µl of cell culture were diluted in series and spotted on LB plates and on 0.5% LB soft agar plates with the susceptible strain PAO1 (one volume of 0.5% LB soft agar at 42 °C, was mixed with 0.01 volume cells at OD 0.5, and poured into a plate). Plates were imaged with the Typhoon 7000 phosphor imager (GE Healthcare) in fluorescence mode.

For *jukA* silencing in PA14, the cells were inoculated 1:100 in MH media with 8 µM final concentration ASOs and every hour 8 µM final concentration ASOs were added additionally until the cells were harvested for CFU/PFU analysis after 3 h.

For PP7 infection, PAO1 cells were inoculated 1:100 in MH media and treated at OD 0.3 with 0.05 µM final concentration ASOs, additionally every 30 min 0.05 µM final concentration ASOs were added. Cells were infected 30 min after starting treatment with PP7 at an MOI of 0.00001, and cells were harvested for CFU/PFU analysis after 2 h.

### Immunoblotting

Infected cell cultures were mixed with final 1×SDS-PAGE loading dye (60 mM Tris/HCl pH 6.8, 0.2 g/ml sodium dodecyl sulphate (SDS), 0.1 mg/ml bromophenol blue, 77 mg/ml DTT, 10% (v/v) glycerol) and were boiled for 10 min at 95 °C for denaturation. Protein samples were analysed by SDS-PAGE and blotted onto methanol-preactivated polyvinylidene (PVDF) membranes. As a loading control, we used Coomassie staining of a second gel where we loaded the same sample volume. ChmA was produced as previously described (Laughlin et al. 2022) in *E. coli* BL21-CodonPlus (DE3)-RIL cells (Agilent Technologies, JVS-12280, chloramphenicol resistance) that were transformed with pET-M14(+) plasmid carrying the *chmA* gene with a N-terminal His-V5-TEV-tag (pMiG118). As previously described ChmA was purified (Laughlin et al. 2022). The tag could not be removed in the purification procedure. Commercial antibody sera were generated at Eurogentec. Rabbits were immunised with the purified protein. The rabbit serum (no. 2481) was used together with anti-rabbit-HRP antibody (Thermo Scientific, 31460) in 5% BSA/TBST for ChmA detection in immunoblotting. Antibody specificity was validated in immunoblotting by comparison between ΦKZ-infected and non-infected cells that yielded a defined band at 70 kDa corresponding for ChmA only in infected cells (**Fig. 1b**).

### Microscopy

0.85% (w/v) agarose was dissolved in 5-fold water diluted LB media and boiled to melt. The liquid agarose was poured on microscope slides with one slide pair at each side as a spacer and one slide on top to form a closed gel slice as described in (Skinner et al. 2013). After solidification ∼1×1 cm pads were cut. Cells were grown in MH media at 220 rpm and 37 °C to an OD600 ∼0.3, then preincubated with ASOs for 30 min, and infected with ΦKZ at an MOI of 5. At indicated time points the phage replication cycle was quenched by cooling the cells on ice for 10 min and the cells were pelleted at 8k×*g* for 5 min. The supernatant was removed, and the cells were resuspended in 500 µl 4% paraformaldehyde and incubated for 15 min on ice. Afterwards cells were washed with PBS and were resuspended in 50 µl PBS for storage at 4 °C. Cells were stained with 16 µM FM4-64, and 360 nM DAPI and 5 µl were layered onto 1.2% agarose pads. The pad was placed with the side of application downwards into a µ-Slide 8 Well high Grid-500. Transmission and fluorescence were detected with a confocal laser scanning microscope Leica SP5. Images were processed with ImageJ (1.53).

For the imaging of ΦKZ155-GFP, PAO1 cells were transformed with a plasmid (pLBu005) coding for ΦKZ155-GFP under the control of an arabinose-inducible pBAD promoter, and selected on gentamycin plates. The cells (JVS-13713) were grown to an OD 0.25 and induced with arabinose at indicated concentrations followed by phage infection with an MOI of 5 at OD 0.3. The harvesting, crosslinking and imaging of cells was conducted as described previously for wt cells.

### RNA preparation and sequencing

PAO1 was grown o/n in LB at 37 °C 220 rpm. Cells were inoculated 1:100 in MH media at 37 °C and grown until OD 0.3. 6 µM ASO (JVPNA-79 for control or -72 for *chmA* inhibition) were added to 1.6 ml of cell culture and incubated for 30 min at 37 °C 220 rpm. Cells were infected with ΦKZ at an MOI of 5. At indicated time points, 250 µl were removed and put on ice. Infection efficiency was independently validated by confocal microscopy and CFU spotting with 50 µl of cells. RNA was isolated from 200 µl of cells using the RNAsnap procedure (Stead et al. 2012). Two volumes of RNAprotect (Qiagen) were added and cells were incubated for 5 min. Cells were pelleted at full-speed for 20 min at 4 °C and the supernatant was removed. The pellet was resuspended in 100 µl SNAP buffer (0.025% SDS, 18 mM EDTA, 1% β-mercaptoethanol, 95% formamide (RNA-grade)). Samples were incubated for 7 min at 95 °C, cell debris was pelleted at full-speed for 5 min at room-temperature, and the supernatant was transferred to a new tube. 1.5 volumes of ethanol were added to the supernatant and the sample was mixed by pipetting. The sample was loaded onto a miRNeasy mini column (Qiagen) two times and spun at full-speed for 20 s at room-temperature. Columns were washed two times with 500 µl RPE buffer (Qiagen) and spun at 8k×*g* for one minute at room-temperature. One final spin was used to dry the column in an empty tube. 30 µl RNase-free water was added to the column and the RNA eluted at 8k×*g* for one minute at room-temperature. The elution was repeated with the flow-through to recover more RNA. The RNA concentration was determined by absorption at 260 nm. RNA was stored at -80 °C.

RNA-sequencing was performed at the CoreUnit SysMed at the University of Würzburg. DNA was digested with DNaseI and the rRNA was depleted with the Lexogen RiboCOP META depletion kit. RNA library was prepared with the CORALL Total RNA-Seq Library Prep Kit V1 (Lexogen). The library was sequenced on the NextSeq2000 (Illumina) with a P1-seq kit (single-end 1x100 bp, Illumina). RNA-seq analysis for the ChmA knockdown and the screen was conducted with READemption 0.4.3 and 2.0.4 (Förstner et al. 2014), respectively. Reads were aligned for PAO1 and ΦKZ to NC_002516 and NC_004629, respectively. Enrichment of transcripts was calculated with the DeSeq2 module in READemption. Read-coverage was illustrated with the Integrated Genomics Viewer (IGV, Robinson et al. 2011).

We defined early and intermediate phage transcripts based on their significant enrichment (log2fold >2 or log2fold <-2, -log10padj >10) between the control 35-and 10-min samples (**Extended Data 2b**, similar as in Ceyssens et al. 2014).

### Structure prediction

Structures were predicted from protein sequences using Google AlphaFold3 server (Abramson et al. 2024, <u>alphafoldserver.com</u>). This information is subject to AlphaFold Server Output Terms of Use found at <u>alphafoldserver.com/output-</u> <u>terms</u> (Google LLC). Ongoing use is subject to AlphaFold Server Output Terms of Use and of any modifications made.

### *In vitro* translation

Template DNA was produced via PCR and Taq-polymerase followed by gel purification. JVO-23244 and -5 were used to amplify wt *ΦKZ155* with a T7 promoter from ΦKZ lysate, and for *ΦKZ155^D102N^* pLBu021 (JVO-) was used as template. The template DNA was amplified with JVO-23244/-5 primers adding T7 promoter to the amplified fragment. 250 ng template DNA was supplemented in 10 µl PURExpress *in vitro* protein synthesis kit mix (NEB). The reaction mix was incubated for 2 h at 30 °C and was subsequently used for assays.

### Cleavage assays

RNA template (JVRNA-001, AUAUAAGGGAACAUAGAUAAACCCCUCCCUAAUAAAAUG) was 5’-^32^P-labelled. For the RNA-DNA duplex, the radiolabelled RNA was mixed in 1:2 ratio together with the reverse complement DNA (JVO-23273), boiled and slowly cooled down to room temperature in a water bath to anneal the duplexes. 1 pmol was added to 2 µl PURExpress IVT mix that translated for 2 h a control, ΦKZ155, ΦKZ155^D102N^, or other nucleases as indicated. The mix was incubated for 1 h, mixed with one volume GLII buffer, boiled for 5 min, rapidly cooled down on ice, and loaded onto 6% 6 M Urea-PAGE (19:1) gels that were run at 300 V for 2 h. The gel was transferred onto Whatman filter paper, vacuum dried, and a phosphor image screen was used to read out the autoradiogram on a Typhoon FLA7000 imager (GE). As a positive control, we used a commercially available RNase H (NEB, M0297S).

### Southern-dot-blotting

PAO1 cells were grown to OD 0.3 in MH media and were pretreated with 6 µM ASOs against ΦKZ nucleases for 30 min. Cells were infected at an MOI of 10 with ΦKZ. At indicated time points a fraction of the culture was removed and 1% SDS was added followed by boiling at 95 °C for 5 min. Subsequently the DNA was extracted from the sample with PCI, and subsequently with one volume chloroform. The aqueous phase was supplemented with 1.5 volume 1 M NaOH and 15 mM EDTA (pH 9). The sample was heated for 3 min at 95 °C and put on ice for 5 min. The solution was filtered with a dot blot apparatus through an equilibrated (0.3 M NaOH) and positively charged nylon membrane. Subsequently, the membrane was dried and the DNA crosslinked via exposition to UV light for 5 min. The membrane was equilibrated with hybridization solution for two times, and a radiolabelled oligo was added for hybridization o/n starting at 60 °C for 1 h and then 48 °C overnight. The membrane was washed once for 15 min with 2×SSC, and 0.5×SSC, and a screen was used for phosphor imaging on a Typhoon FLA7000 imager (GE).

### Data availability

Raw sequencing data and coverage files are accessible at Gene Expression Omnibus (Barrett et al., 2012) for ChmA knockdown and the screen experiments with the accession numbers GSE269401 and GSE269911, respectively. Analysed data are listed in **Suppl. Data 2 and 3**.

## Acknowledgements

We thank Laura Vogel and Barbara Plaschke for technical assistance, and Esther Hauschild for microscopy. We thank Anke Sparmann for discussions and for editing the manuscript. We thank the Core Unit SysMed at the University of Würzburg and Tom Gräfenhahn for excellent technical support and RNA sequencing. The work was funded by Deutsche Forschungsgemeinschaft (DFG) project 465133664 in the Priority Programme “New Concepts in Prokaryotic Virus-host Interactions – From Single Cells to Microbial Communities” (SPP 2330) awarded to J.V.. This work was also supported by a Gottfried Wilhelm Leibniz award (DFG Vo875/ 18) awarded to J.V..

